# Seoul orthohantavirus evades innate immune activation by reservoir endothelial cells

**DOI:** 10.1101/2024.06.18.599607

**Authors:** Stefan Klimaj, Autumn LaPointe, Kimberly Martinez, Eduardo Hernandez Acosta, Alison M. Kell

## Abstract

Pathogenic hantaviruses are maintained world-wide within wild, asymptomatic rodent reservoir hosts, with increasingly frequent human spillover infections resulting in severe hemorrhagic fever disease. With no approved therapeutics or vaccines, research has, until recently, focused on understanding the drivers of immune-mediated pathogenesis. An emerging body of work is now investigating the mechanisms that allow for asymptomatic, persistent infections of mammalian reservoir hosts with highly pathogenic RNA viruses. Despite limited experimental data, several hypotheses have arisen to explain limited or absent disease pathology in reservoir hosts. In this study, we directly tested two leading hypotheses: 1) that reservoir host cells induce a generally muted response to viral insults, and 2) that these viruses employ host-specific mechanisms of innate antiviral antagonism to limit immune activation in reservoir cells. We demonstrate that, in contrast to human endothelial cells which mount a robust antiviral and inflammatory response to pathogenic hantaviruses, primary Norway rat endothelial cells do not induce antiviral gene expression in response to infection with their endemic hantavirus, Seoul orthohantavirus (SEOV). Reservoir rat cells do, however, induce strong innate immune responses to exogenous stimulatory RNAs, type I interferon, and infection with Hantaan virus, a closely related hantavirus for which the rat is not a natural reservoir. We also find that SEOV-infected rat endothelial cells remain competent for immune activation induced by exogenous stimuli or subsequent viral infection. Importantly, these findings support an alternative model for asymptomatic persistence within hantavirus reservoir hosts: that efficient viral replication within reservoir host cells prevents the exposure of critical motifs for cellular antiviral recognition and thus limits immune activation that would otherwise result in viral clearance and/or immune-mediated disease. Defining the mechanisms that allow for infection tolerance and persistence within reservoir hosts will reveal novel strategies for viral countermeasures and inform rational surveillance programs.

**Author Summary:** Despite the significant, and continual, threat to human health, limited experimental evidence explains the mechanisms that underly asymptomatic zoonotic RNA virus persistence within natural, mammalian reservoir hosts. Here, we investigated whether reservoir host target cells for hantavirus infection are competent for antiviral activation and tested the hypothesis that, through long-term co-evolution, Seoul orthohantavirus actively antagonizes innate antiviral signaling pathways to limit immune induction and prevent viral clearance in primary reservoir cells. While we find no evidence to support these hypotheses, our findings do support an alternative hypothesis that viral replication within the natural reservoir cells may not result in the production of immune-stimulating by-products. The factors that determine viral persistence within the reservoirs may include efficient use of host co-factors for efficient genome replication and/or packaging and warrant further investigation.

## Introduction

Zoonotic RNA viruses pose ever-present threats to human health. Maintained within natural reservoir host populations, they have the potential for human spillover and disease with rising frequency due to intensifying habitat encroachment and climate change. Viral dynamics within natural reservoir populations provide ecological clues for viral maintenance that can be applied to calculate the threat of zoonotic events (1–4). Often zoonotic RNA viruses cause little-to-no disease within natural reservoirs, even in the presence of high viral loads (5–10). Two leading hypotheses to explain host-specific infection outcomes include reservoir host disease tolerance due to heightened or dampened general immune responses and co-evolved virus-host interactions that antagonize immune signaling pathways (11–13). In fact, different viral species likely employ unique mechanisms to modulate reservoir host responses toward the goal of persistence.

Defining these mechanisms for mammalian reservoir hosts against viruses that otherwise cause severe disease in humans may shed light on the factors that drive pathogenesis, limit viral replication, and facilitate viral maintenance within reservoir populations.

Viral detection and the initiation of type I and type III interferon (IFN) responses within infected cells are critical to initiate an effective antiviral response capable of restricting viral replication, limiting further tissue dissemination, and facilitating viral clearance (14). Double-stranded RNAs (dsRNA), highly structured RNAs, and unprocessed RNAs, which are recognized by the cytosolic RIG-I-like receptors (RLR), retinoic acid-induced gene I (RIG-I) melanoma differentiation-associated protein 5 (MDA5), drive transcription of antiviral effectors and types I and III IFNs (14–17). The critical importance of this pathway in constraining viral replication and transmission is evidenced by almost universal antagonism of type I IFN signaling or effector function by successful viral pathogens (17). Notably, while the intended outcome of type I IFN induction is viral clearance, strong and sustained IFN responses are implicated in inflammatory disease, cancers, and significant tissue damage following viral infection (18–22). Tight regulation of these pathways by the host allows for efficient antiviral response with limited tissue damage. Similar selective pressures also apply to viruses for which it would be beneficial to limit host immune activation to evade restriction and establish a replicative niche for long-term persistence and transmission.

The family Hantaviridae (Order *Bunyavirales*) is composed of insectivore- and rodent-borne viruses, maintained within reservoir populations through direct transmission (23–25). Human infection with rodent-borne orthohantaviruses (here referred to as hantaviruses) occurs through inhalation of excreta from infected reservoirs, with endothelial cells (EC) being the primary cellular target for replication. Currently, no FDA or EU-approved vaccines or therapeutics exist that specifically target hantaviruses, despite case fatality rates of up to 60%. Severe human hantavirus disease is characterized by high levels of circulating proinflammatory factors (IL-6, TNFα), thrombocytopenia, and vascular leakage (26–29). Hantaviruses are non-cytopathic and, in fact, inhibit both intrinsic and extrinsic apoptotic pathways in human EC (30–32). Therefore, this robust inflammatory response to hantavirus infection is thought to be a central driver of tissue damage and severe human disease (33). We previously reported that the RLRs are essential for interferon stimulated gene (ISG) expression in Hantaan orthohantavirus (HTNV)-infected human umbilical vein endothelial cells (HUVEC) (34). In wild-type HUVEC, we observed robust ISG expression beginning 48hrs post-infection, which was delayed in RIG-I^-/-^ cells and ablated in the absence of both RLR. We further found that UV-inactivated HTNV did not drive ISG induction, supporting the hypothesis that viral replication intermediates are the primary pathogen-associated molecular patterns (PAMPs) for hantaviruses in human cells. Importantly, humans are a dead-end host for hantaviruses, with extremely rare exceptions for Andes orthohantavirus, and the antiviral response appears to be effective for viral clearance in survivors and fatal victims of hantavirus infection (35–38). Whether sustained activation of type I IFN responses in human EC serves as the catalyst of an unchecked systemic inflammatory syndrome and severe hantavirus disease, despite apparent viral control, remains to be explored.

In contrast, *in vivo* hantavirus infection in natural reservoir hosts leads to a very mild systemic antiviral response early, followed by a dominant regulatory T cell signature, despite high viral loads (39–42). Importantly, individual hantavirus species are closely associated with specific mammalian reservoir hosts, leading to a hypothesis of extended co-evolution between virus and reservoir host (25, 43). For example, Sin Nombre orthohantavirus is found almost exclusively within *Peromyscus* (Deer mouse) species in the Americas, HTNV has been isolated from *Apodemus agrarius* (striped field mouse) in Asia, and Seoul orthohantavirus (*Orthohantavirus seoulense*, SEOV) has been identified in *Rattus* species (predominantly Norway rat) throughout the world (44–49). Previous work by several groups has established a paradigm in which infection of natural hantavirus reservoir rodents with their endemic, potentially co-evolved, virus species results in asymptomatic, persistent infection, whereas infection of that rodent species with a non-endemic hantavirus leads to rapid antiviral activation and viral clearance (50–53). *In vitro* infections of reservoir endothelial cells with their endemic hantavirus do not stimulate antiviral gene expression mount a mild, if any, innate immune response to their endemic hantavirus (54, 55). However, the mechanisms that underlie immune recognition and infection outcome for each unique virus-host relationship are poorly defined.

Although tools to identify novel zoonotic viruses and to systematically document reservoir prevalence have rapidly improved in recent decades, the virus-host interactions that allow for asymptomatic virus persistence within these natural hosts remain elusive. Several hypotheses to explain persistence within reservoir hosts have been put forth, with a few becoming widely accepted, despite limited experimental support (11–13). The co-evolved nature of human pathogenic hantaviruses with their reservoir rodent hosts provides us with a unique opportunity to test these hypotheses at the cellular level. Here, we describe robust innate immune activation and inflammatory responses in human EC in response to SEOV infection that are dependent on RIG-I-like receptor activity. In contrast, we show that primary reservoir rat EC do not mount an antiviral ISG response to their endemic hantavirus, SEOV, despite robust infection and replication. We show that RLMVEC upregulate ISGs in response to non-endemic hantavirus infection, type I IFN, and known RLR agonists. We rigorously tested the hypothesis that SEOV directly antagonizes innate immune signaling pathways by interfering with RLR or type I IFN signaling and found no evidence for it. The studies presented here instead suggest a more complex hypothesis: that endemic hantaviruses may replicate more efficiently within reservoir target cells to prevent immune recognition and antiviral activation. These studies add to an emerging body of knowledge to better understand the unique mechanisms that allow for persistence of highly pathogenic zoonotic viruses within reservoir host populations. Defining the mechanisms employed by zoonotic viruses and their hosts to regulate immune-mediated damage induced by infection will reveal novel strategies for therapeutic development against severe human disease.

## Results

### Reservoir and human EC are differentially susceptible to SEOV infection

It has been previously shown that both HUVEC and RLMVEC are susceptible to SEOV infection, however a quantitative assessment of SEOV infection in comparison to immune-incompetent Vero E6 cells has not been reported. Because hantaviruses are non-cytopathic, a focus-forming unit assay, reliant on antibody staining for the viral nucleocapsid (N) protein, in Vero E6 cells is the established method for titering virus stocks (56). However, primary cells are less tolerant of the seven day incubation period and methylcellulose overlay required for this assay, making it challenging to determine viral titer on relevant, immune-competent cells. We therefore developed a novel protocol for tittering our viral stocks on Vero E6, RLMVEC, and HUVEC. We infected cells with SEOV (strain SR11) at 2-fold dilutions (from undiluted to 1:1024) for one hour followed by the addition of a 2% methylcellulose overlay. We then fixed the cells 24 hours post-infection and probed with a highly specific monoclonal antibody against SEOV N, fluorescently-labeled secondary antibody, and DAPI to stain the nuclei (**Fig 1a**). Using the Thermo Scientific CellInsight CX7 High-content screening platform we quantified the percentage of N+ cells at each dilution (**Fig 1b**). A similar methodology to quantify Puumala and Tula orthohantavirus infected cells using flow cytometry was recently reported (57). Menke and colleagues proposed a simple equation to determine infectious unit count/milliliter (mL) by multiplying the fraction of infected cells by total cell number and dilution (e.g. 1:10). Applying this equation to our quantification results derived from microscopy, we were able to quantify the infectious units/mL in our virus stock (**Fig 1c**). We calculated a similar number of infectious units/mL on Vero E6 cells using both the traditional FFU assay (6x10^5^) and our novel microscopy assay (2x10^6^). Importantly, we determined that the number of infectious units within the same virus stock in RLMVEC was only slightly higher than that for Vero E6 cells, suggesting that both cell types are equally susceptible to virus infection. Conversely, the number of infectious units/mL quantified for HUVEC was more than a log lower than Vero E6 or RLMVEC, indicating that these cells are less susceptible to SEOV infection. We therefore calculated viral titers for our SEOV stocks for each of our cell types of interest and report here the multiplicity of infection (MOI) in subsequent experiments based on these cell-type specific titers.

**Figure 1.**
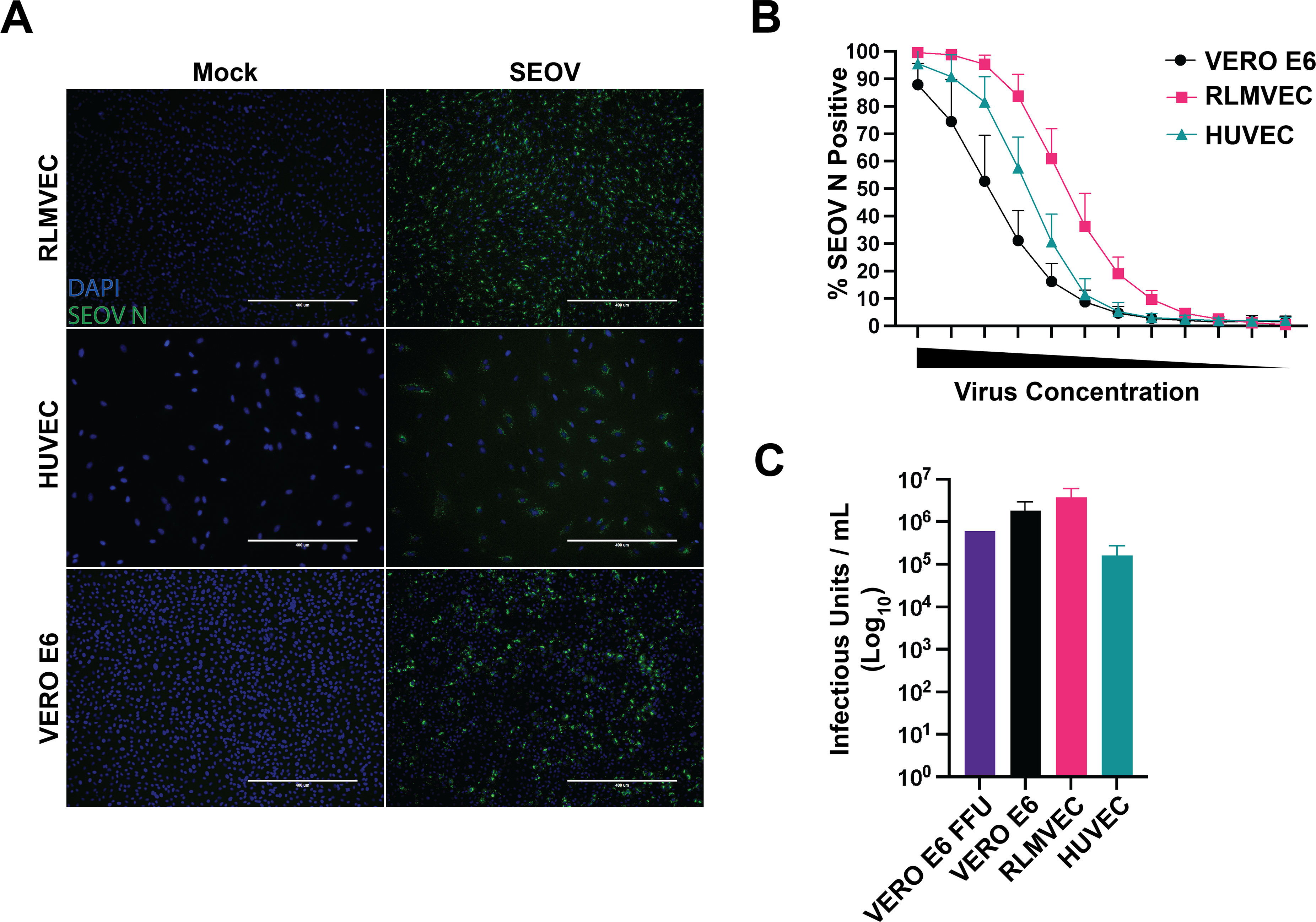
Cell-type specific titers in human and rat EC. Primary RLMVEC, HUVEC, and Vero E6 cells were infected with serial dilutions of SEOV to quantify the number infectious units present in viral stocks for each cell type. A) Images of mock or SEOV-infected (1:8 dilution) RLMVEC, HUVEC, or Vero E6 cells at 10x magnification on EVOS FL Auto Imaging system. Nuclei stained with DAPI (blue) and cells probed for SEOV N (green). B) Percentage of cells determined to be positive for SEOV N expression calculated using the CellInsight CX7 imaging platform for each cell type following infection at decreasing 1:2 dilutions of viral stock. C) Calculated titer for each cell type based on either FFU (Vero E6 only) or CellInsight CX7 imaging calculations (± SD). Summary data from four replicate wells per dilution performed in three independent experiments.

### Seoul virus infection drives RIG-I-dependent ISG expression in HUVEC

To determine whether the RLRs are required for recognition of, and immune signaling against, SEOV, we infected HUVEC lacking either RIG-I or MDA5, or both RLRs at MOI 0.01. Similar to our HTNV observations, SEOV infection in Cas9 control cells led to increased ISG expression over time, concomitant with SEOV nucleocapsid detection (**Fig 2a**). We observed that RIG-I^-/-^ and RLR^-/-^ HUVEC lacked all ISG induction following SEOV infection, with MDA5^-/-^ resembling the wild-type control. Therefore, RLR-dependent innate immune activation may be a conserved human EC response to Old World hantavirus infections, with HTNV activating both RIG-I and MDA5 and RIG-I being the critical PRR for SEOV. We next asked whether SEOV infection induces inflammatory cytokine and chemokine signaling in an RLR-dependent fashion. WT HUVEC or RLR^-/-^ cells were infected with SEOV and supernatant was interrogated for secreted cytokines and chemokines four days post-infection (**Fig 2b**). As expected, we observed an increase in neutrophil, monocyte, and NK cell recruitment factors CXCL10, CCL5, and IL-15 during SEOV infection in Cas9 WT control HUVEC. This induction was lost in SEOV-infected RLR^-/-^ HUVEC. These results suggest that RLR signaling in response to SEOV infection in human EC may also catalyze a larger proinflammatory state within the vasculature through chemokine secretion and endothelial activation.

**Figure 2.**
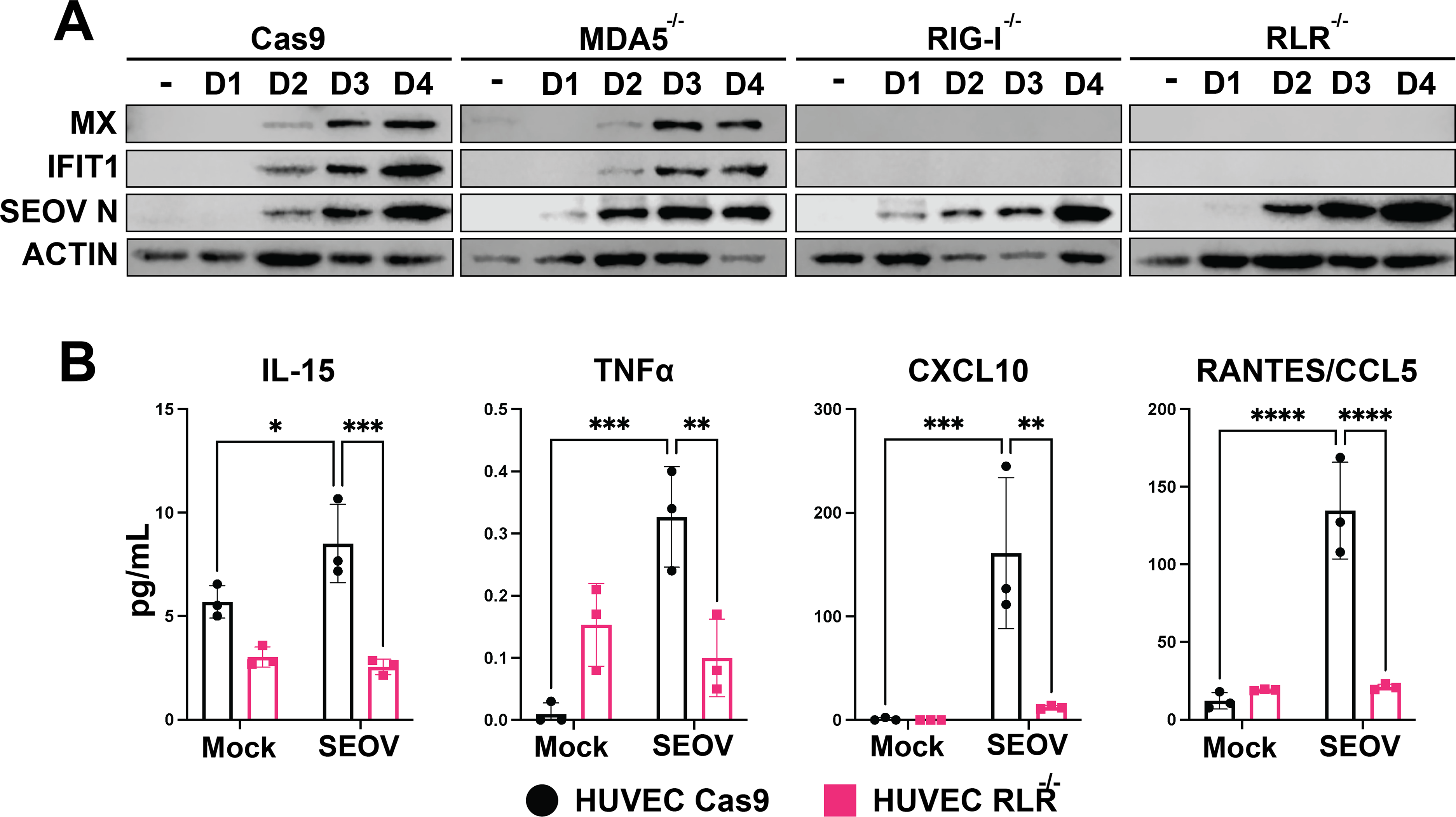
SEOV-induced ISG expression in HUVEC is RIG-I-dependent. CRISPR-modified HUVEC (Cas9 alone, RIG-I^-/-^, MDA5^-/-^, or both RIG-I-like receptors (RLR^-/-^) were infected with SEOV at MOI 0.01. Cell lysates and supernatant were collected every 24 hours for four days. A) Cell lysates were run on SDS-PAGE and subjected to immunoblot analysis. Data shown are representative of three independent experiments. B) Supernatants of SEOV-infected cells were collected on day four and analyzed for cytokine and chemokine expression by Eve Technologies. Data represent three biological replicates ± SD. Two-way ANOVA with multiple comparisons, * denotes adjusted p<0.05.

### Seoul virus infection in primary rat EC does not drive ISG expression

Next, we interrogated reservoir target cells for antiviral responses to endogenous and non-endogenous hantavirus infection. We infected RLMVEC with either SEOV (endogenous hantavirus) or HTNV (closely related but for which rats are not the reservoir host) at 0.05 MOI **(Fig 3a)**. While HTNV-infected cells increased expression of the ISGs RIG-I, MDA5, and Mx1/2/3 over the course of the six-day infection, SEOV did not drive increased ISG production. Similar results were observed at the transcript level on day two post-infection for antiviral genes *Ifit3*, *Mx1*, and *Oas1* **(Fig 3b)**. Importantly, as we have observed previously in HUVEC, UV-inactivated HTNV did not induce ISG expression in these cells, indicating that antiviral signaling requires viral replication **(Fig 3c)** (58). To confirm that these results are not strain-specific, we infected primary RLMVEC with the Baltimore strain of SEOV with consistent observations (**Fig 3d**) (59). Previous reports have indicated a potential loss of pathogenicity for hantaviruses following culture in Vero E6 cells (60, 61). Therefore, we passaged our SR-11 strain of SEOV exclusively in RLMVEC three times and then assessed ISG expression in RLMVEC (**Fig 3e**). Importantly, we again observed no induction of ISGs following infection of RLMVEC with these SEOV variants, but strong induction when infected with HTNV. UV-inactivated SEOV was included as a control to determine whether cytokines expressed during propagation on RLMVEC could drive ISG expression, although this was not observed. These results demonstrate that primary rat cells induce antiviral responses to a non-endemic hantavirus, HTNV, but fail to respond to SEOV infection despite robust replication.

**Figure 3.**
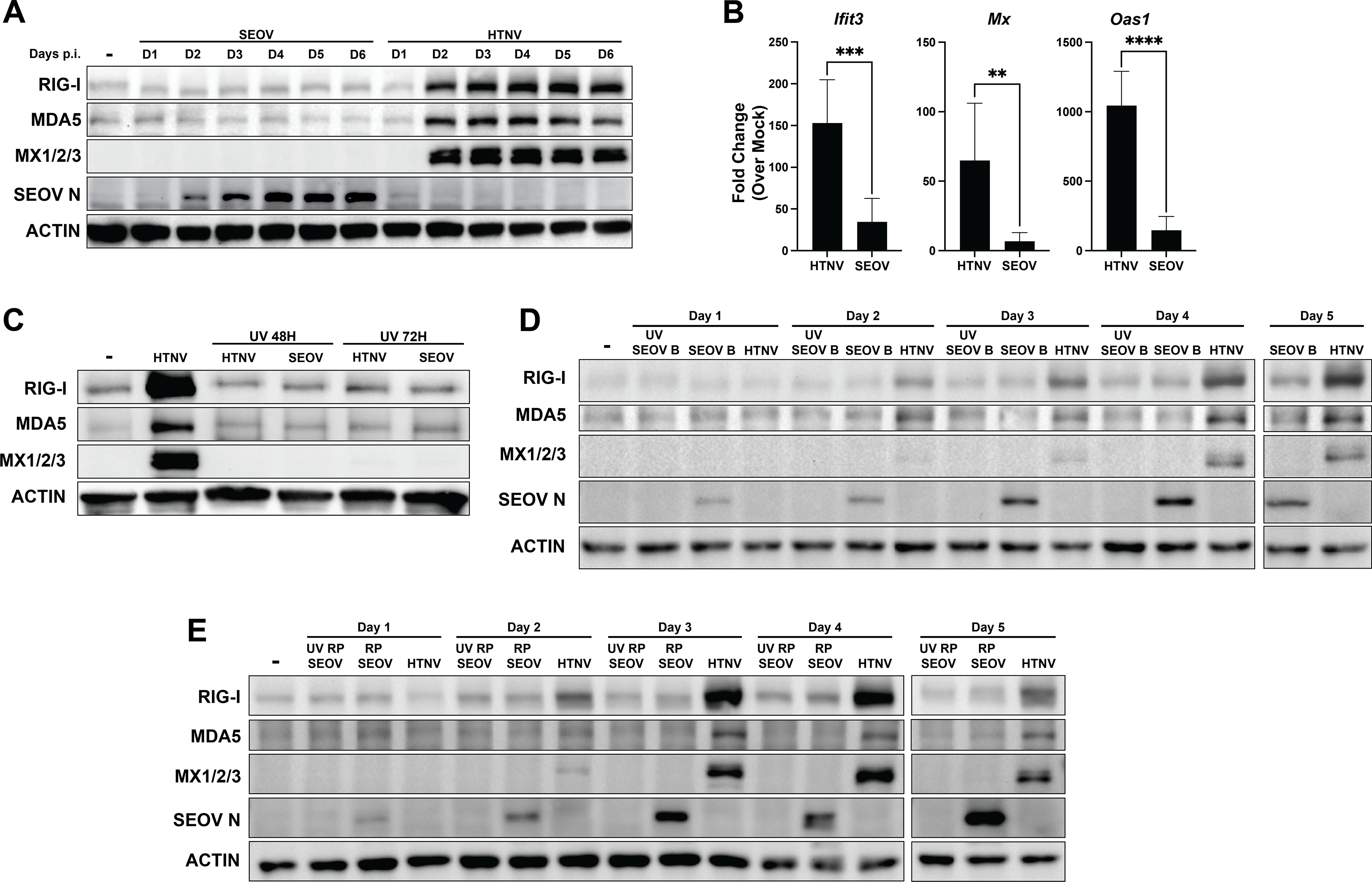
Primary rat EC induce ISGs in response to non-endemic hantavirus, HTNV, but not to endemic hantavirus, SEOV. A) RLMVEC were infected with either SEOV or HTNV (MOI 0.05) or mock infected (-) and lysates were collected every 24 hours for six days (D1, D2, etc.) post-infection. Cell lysates were subjected to SDS-PAGE and immunoblot analysis for ISG expression. B) RLMVEC were infected with either SEOV or HTNV (MOI 0.05) and harvested 48 hours post-infection. Gene expression was assayed by comparative RT-PCR and averaged data from three experimental replicates is shown. Student’s T test, * denotes p<0.05. C) RLMVEC were infected with either HTNV or UV-inactivated HTNV or SEOV (MOI 0.05). Lysates were harvested at the indicated times post-infection and subjected to immunoblot analysis. D) RLMVEC were infected with either SEOV Baltimore strain (SEOV B), UV-inactivated SEOV Baltimore strain (UV SEOV B) or HTNV (MOI 0.05). Lysates were harvested at the indicated times post-infection and subjected to immunoblot analysis. E) RLMVEC were infected with either RLMVEC-passaged SEOV (RP SEOV), UV-inactivated RLMVEC-passaged SEOV (UV RP SEOV), or HTNV (MOI 0.025) and lysates were harvested at the indicated times post-infection and subjected to immunoblot analysis. All data shown are representative of ≥3 independent experiments.

### Primary rat EC maintain intact antiviral responses to exogenous RNA stimuli

Altered immune function or tolerance is a popular explanation for the seemingly unique ability of rodent and bat reservoir hosts to support significant and diverse pathogen burden (5–7, 9). Therefore, we investigated the ability of primary RLMVEC to induce antiviral gene expression following treatment with known stimulatory RNA and exogenous type I interferon. Rat ECs transfected with poly(I:C), a dsRNA mimic which activates RLR-dependent signaling, or treated with recombinant rat IFNβ, increased relative transcription of *Cxcl10* and *Ccl2* compared to mock-treated controls **(Fig 4a).** X RNA is an *in vitro* transcribed 100-nucleotide sequence derived from the hepatitis C virus genome that does not drive RLR activation and serves as an additional control (62, 63). Increased expression at the protein level for RIG-I, MDA5, and Mx1/2/3 was also observed for RLMVEC treated with exogenous IFNβ and transfected poly(I:C) (**Fig 4b)**. Curiously, we did not detect ISG expression following addition of poly(I:C) to the cell culture media, which has been reported to stimulate TLR3 signaling (64). We next asked whether RNA PAMPs created during SEOV infection could stimulate an antiviral response in the *absence* of viral proteins. To test this, we isolated total RNA from SEOV-infected RLMVEC five days post-infection, capturing all host and viral RNAs present within a robustly infected culture. Total RNA from uninfected RLMVEC (cRNA) was also isolated as a negative control. We then transfected the infected-cell RNA (icRNA) into uninfected RLMVEC and measured relative antiviral gene expression 18 hours later. We observed a dose-dependent increase in ISG transcription following transfection of icRNA when compared to cells transfected with cRNA **(Fig 4c)**. Together, these data demonstrate that primary RLMVEC are immune-competent. Further, SEOV RNA replication intermediates, or other viral-induced host RNAs, stimulate an innate immune response in reservoir EC in the absence of viral proteins.

**Figure 4.**
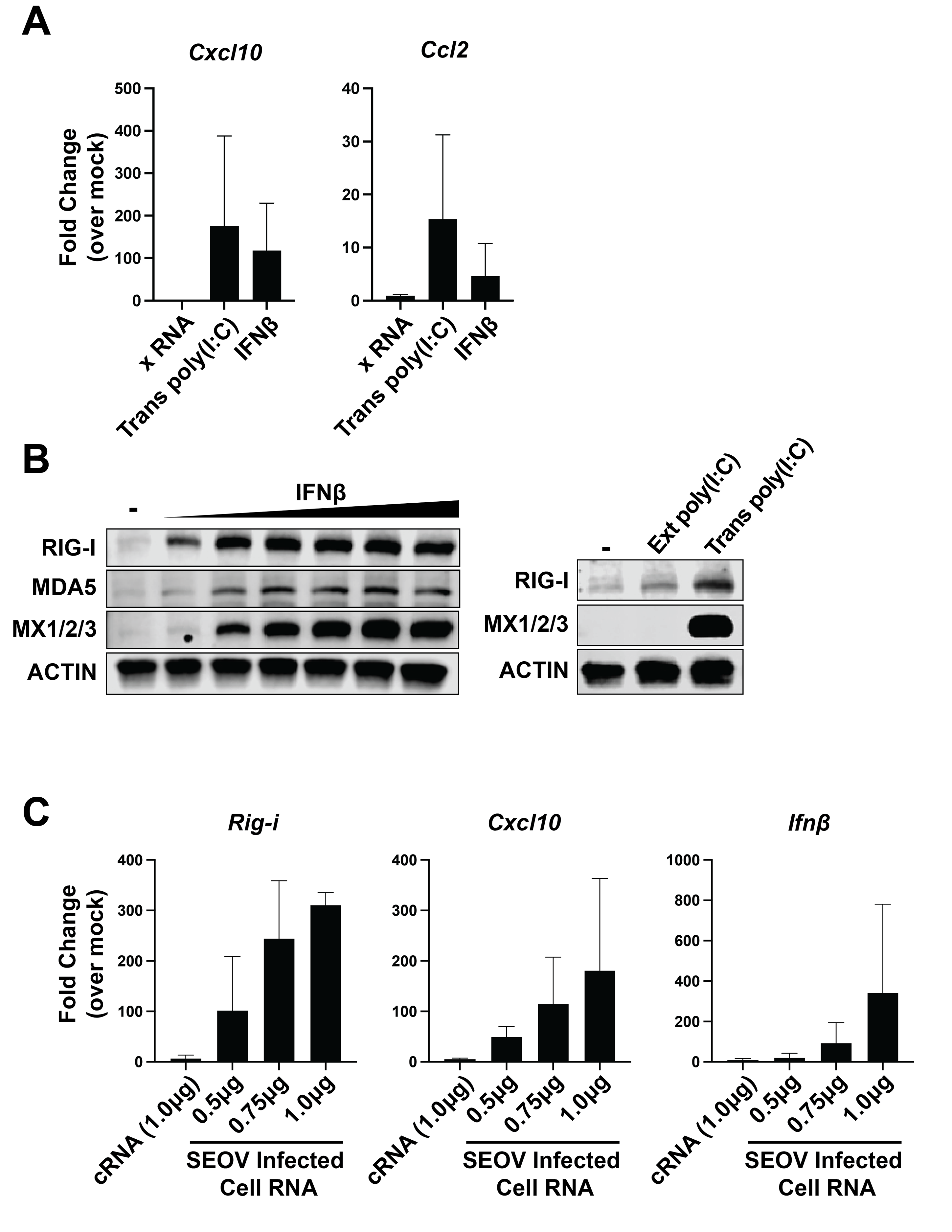
Primary RLMVEC induce antiviral responses to canonical signaling agonists. A, B) RLMVEC were either treated with exogenous recombinant rat IFNβ (1-100U/mL) or transfected with HCV-derived X RNA or poly(I:C) (100ng). RNA (18 hours) and lysates (24 hours) were collected post-treatment and subjected to comparative RT-PCR analysis (A) or immunoblotting (B). C) RLMVEC were transfected with indicated amounts of total RNA isolated from either mock-infected Vero E6 cells (cRNA) or SEOV-infected Vero E6 cells (icRNA). RLMVEC RNA was collected 18 hours post-transfection for comparative RT-PCR analysis. All data shown represent ≥3 independent experiments, ±SD.

### Active SEOV infection does not antagonize antiviral gene expression in primary rat EC

Because immune-competent RLMVEC respond to viral and/or viral-induced RNA in the absence of viral proteins but do not induce ISGs during active SEOV infection, we hypothesized that SEOV proteins may inhibit RLR signaling or autocrine type I IFN activation in reservoir rat cells. To test this hypothesis, we infected RLMVEC with SEOV at MOI 0.05 and allowed the virus to replicate and produce viral proteins for 48 hours. We then either treated cells with transfected poly(I:C) (**Fig 5a**), recombinant rat IFNβ (**Fig 5b**), or superinfected with HTNV (**Fig 5c**). If SEOV infection actively antagonizes either RLR or type I IFN signaling cascades, we would expect a reduction in ISG expression in cells that were first infected with SEOV compared to uninfected cells treated with innate immune agonists alone. We observed no such decrease in gene expression in SEOV-infected poly(I:C)-transfected cells when compared to poly(I:C) transfection alone (**Fig 5a**). Similarly, no change in gene expression was observed in response to IFNβ treatment with or without prior SEOV infection (**Fig 5b**). Finally, HTNV also induced robust ISG protein expression in primary RLMVEC, irrespective of SEOV infection status (**Fig 5c**). Our previous studies of HTNV immune induction in human EC revealed a dependence on the RLR for ISG expression. We, therefore, generated RLMVEC that lack either RIG-I or MDA5 protein expression to test whether prior SEOV infection could block recognition of HTNV and ISG expression through a single RLR. For example, if SEOV specifically counteracts the activation of rat RIG-I, we would expect that HTNV would fail to induce ISG expression in SEOV-infected MDA5^-/-^ RLMVEC, due to a loss in functionality of both RLRs. In **Figure 5d**, we show that, as expected, the lack of either RIG-I or MDA5 reduces the expression of Mx1/2/3 following HTNV infection. However, we observe no difference in ISG expression between cells infected with HTNV alone compared to cells first infected with SEOV and then HTNV in either WT or KO cells. Finally, to confirm that HTNV-driven ISG expression was in fact occurring in SEOV-infected cells, we performed immunofluorescence assays probing for Mx1/2/3 protein expression 48hrs post-HTNV superinfection (**Fig 6**). By co-staining for SEOV nucleocapsid and Mx1/2/3, we observed that Mx signal was produced in SEOV N (+) cells (white arrows). Regrettably, a lack of specific antibodies against HTNV nucleocapsid prevents identification of HTNV-infected cells in this assay. However, this result demonstrates that SEOV infection does not limit the induction and signaling of type I IFN. We therefore conclude that direct antagonism of either RLR- or type I IFN-signaling pathways is not the mechanism by which SEOV is able to replicate in primary reservoir EC without triggering robust immune activation.

**Figure 5.**
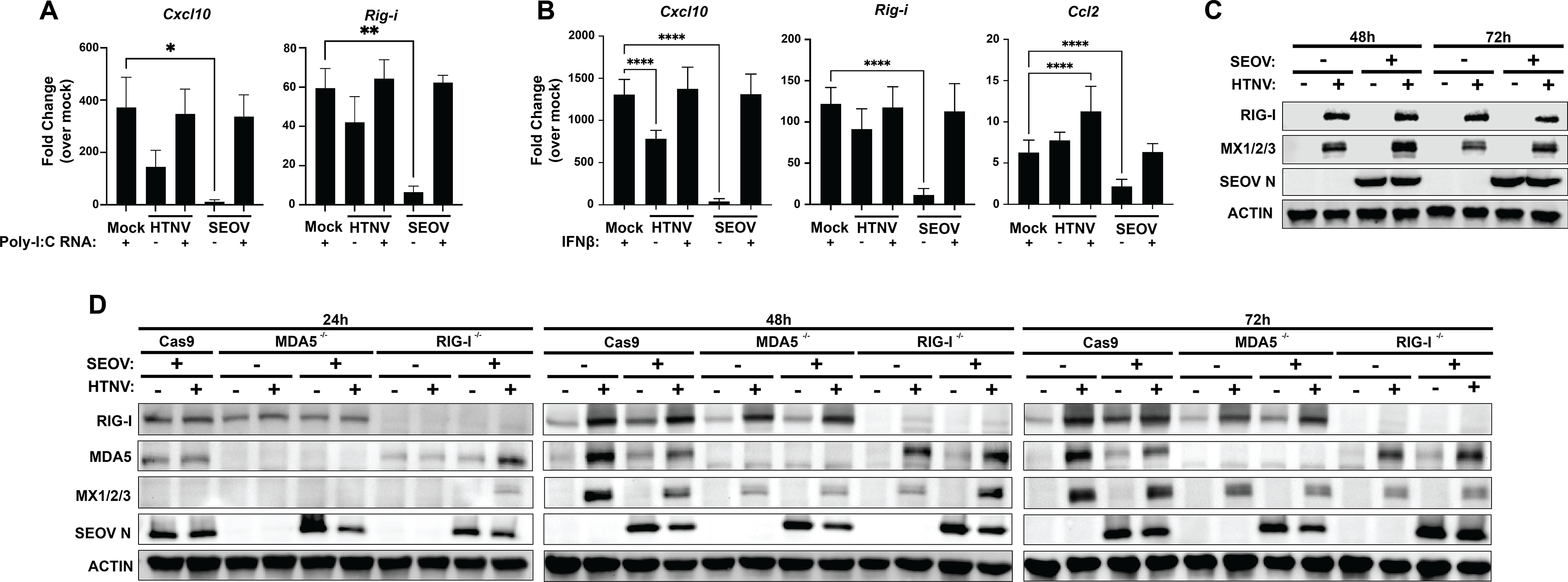
SEOV infection does not inhibit subsequent ISG induction. A-B) RLMVEC were mock-infected or infected with either SEOV or HTNV (MOI 0.1) for 24 hours and then (A) transfected with poly(I:C) (100ng) or (B) treated with IFNβ (100U/mL). RNA was collected 24 hours following poly(I:C) or IFNβ treatment. Gene expression was assayed by comparative RT-PCR. One way ANOVA with multiple comparisons to mock-infected poly(I:C)/IFNβ control, * denotes p<0.05. C) RLMVEC were either mock-infected or infected with SEOV (MOI 0.05) 48 hours. Cells were then either subsequently superinfected with HTNV (MOI 0.05) or mock-infected. Cell lysates were collected 48 hours and 72 hours post-HTNV infection and subjected to SDS-PAGE and immunoblot analysis. D) RLMVEC Cas9 scramble, RIG-I^-/-^, or MDA5^-/-^ cells were either mock-infected or infected with SEOV (MOI 0.05) 48 hours. Cells were the either subsequently superinfected with HTNV (MOI 0.05) or mock-infected. Lysates were collected at the indicated times post-HTNV infection and subjected to SDS-PAGE and immunoblot analysis. All data shown represent ≥3 independent experiments, ±SD.

**Figure 6.**
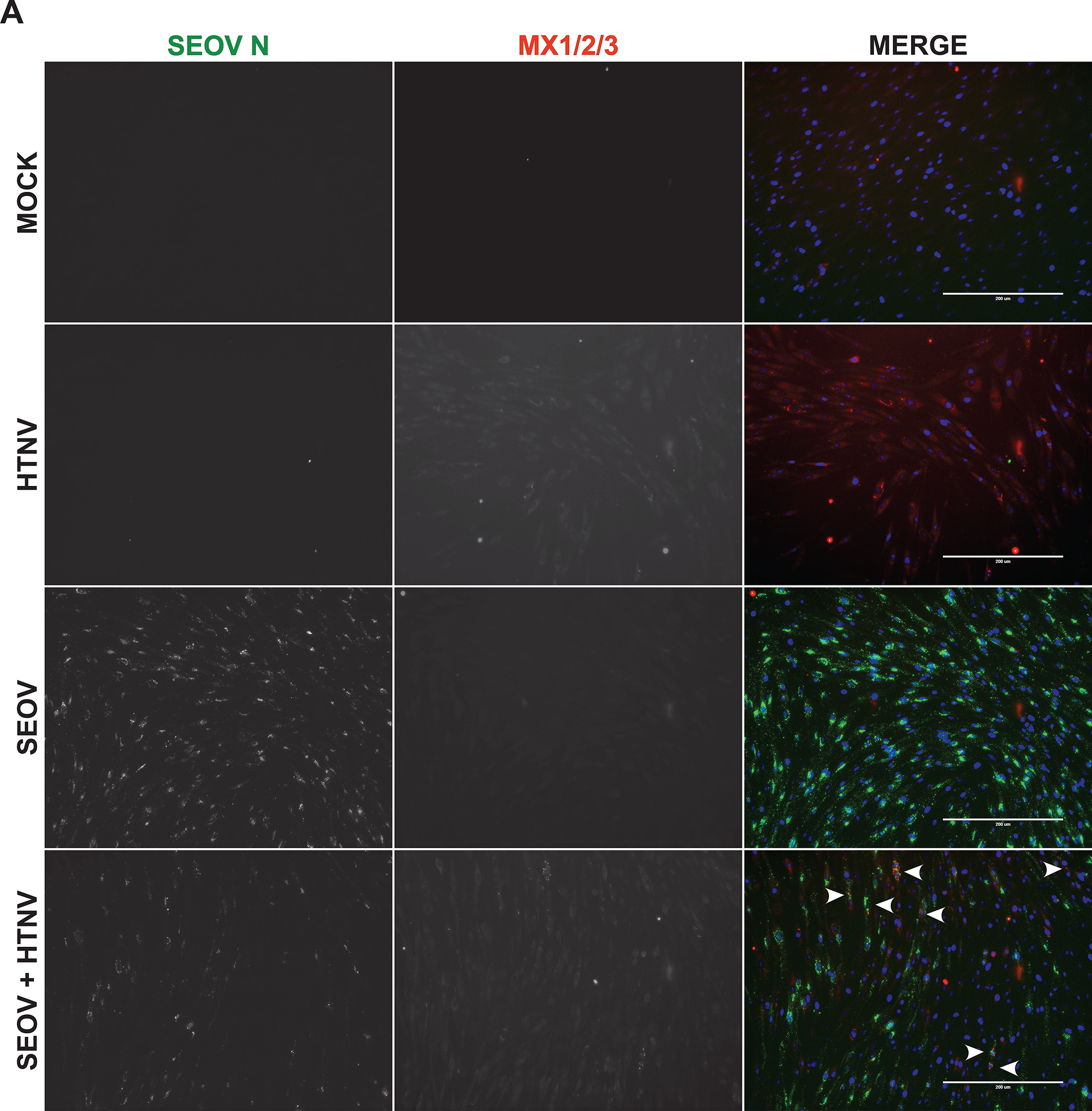
SEOV does not inhibit HTNV-induced Mx1/2/3 expression. RLMVEC were either mock-infected or infected with SEOV (MOI 0.05) for 48 hours. Cells were subsequently either mock-infected or superinfected with HTNV (0.05 MOI) and fixed 48 hours post-HTNV infection. Images of immunostaining for SEOV nucleocapsid (green), MX1/2/3 (red), and DAPI (blue). Images were captured on an EVOS FL Auto imaging system using 20x magnification. Arrows indicate individual cells expressing both Mx1/2/3 and SEOV N. Images are representative of 2 independent experiments.

### Increasing SEOV MOI leads to RLR-dependent innate immune signaling in RLMVEC

Given these results, and the robust susceptibility of RLMVEC to SEOV infection, we tested whether increasing viral dose might drive antiviral activation in reservoir cells. Western blot analysis revealed a dose-dependent increase in ISG expression with increasing MOI (**Fig 7a**). Notably, viral nucleoprotein is detected in all infection conditions, revealing that the number of incoming infectious units at the initial infection of the cell culture is what determines immune activation in these reservoir cells. We, therefore, hypothesized that high MOI infections may lead to increased production of defective viral genomes or accumulation of defective replication intermediates that may be recognized by RLRs to induce antiviral responses (65–68). To test this hypothesis, we infected our CRISPR knockout primary RLMVEC, lacking RIG-I or MDA5, at a range of MOI. Across several experiments we determined that either RIG-I or MDA5 were exclusively required for ISG expression (**Fig 7b**). This inconsistency between experiments suggests that the RLRs may have somewhat redundant roles for signaling in response to high MOI SEOV infection in rat EC. These findings again demonstrate that primary RLMVEC are competent for sensing viral infection, but that low MOI SEOV infections allow the virus to subvert antiviral activation despite productive spread in culture.

**Figure 7.**
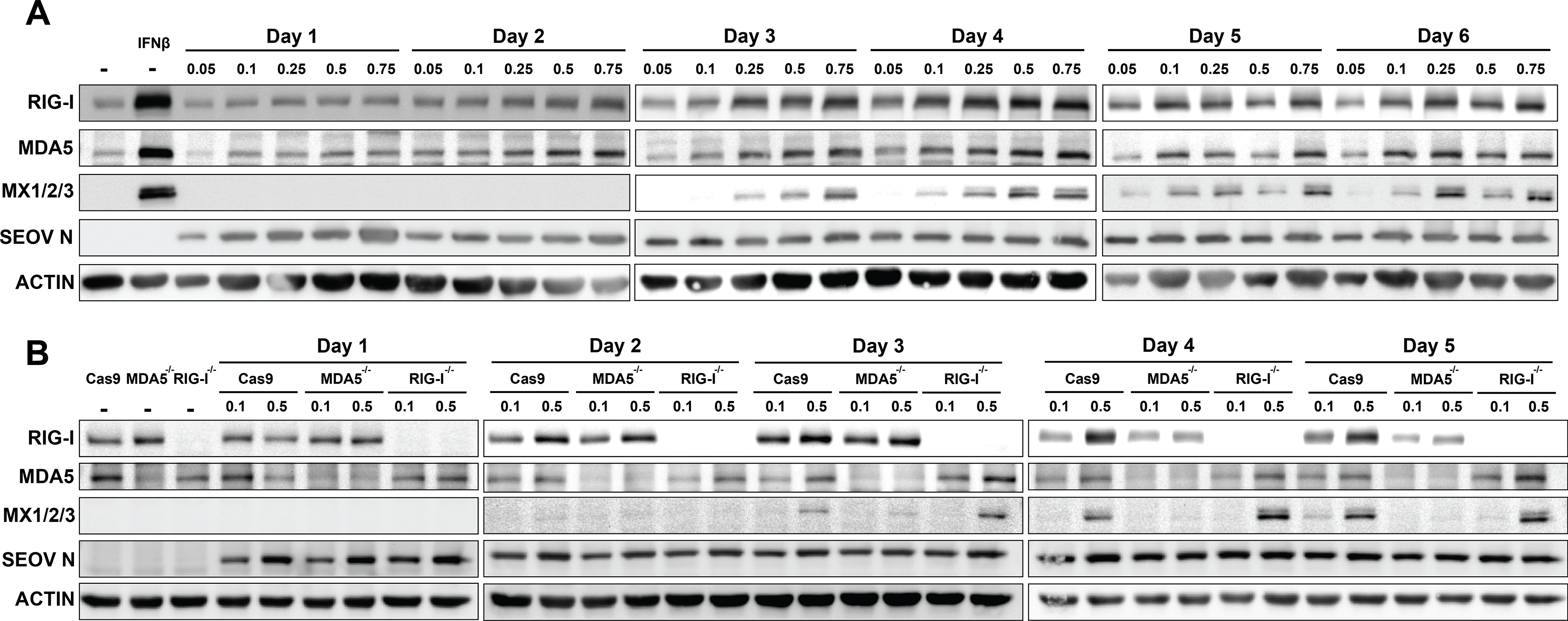
High MOI infections drive RLR-dependent ISG expression in RLMVEC. A) RLMVEC were infected with SEOV at the noted MOI 0.05 – 0.75 and cell lysates collected every 24 hours as indicated. Cell lysates were subjected to immunoblot analysis. Mock-infected RLMVEC (-) treated with 100U/mL IFNβ 24 hours prior to harvest serves as a positive control. B) Cas9 scramble, RIG-I^-/-^, and MDA5^-/-^ RLMVEC were mock-infected (-) or infected with SEOV at either MOI 0.1 or 0.5 and harvested at the indicated times post-infection. Lysates were subjected to immunoblot analysis. All data shown represent ≥3 independent experiments.

## Discussion

Our study investigated the potential cellular mechanisms underlying hantavirus persistence within their asymptomatic, natural reservoir hosts. We have presented evidence that primary rat reservoir target cells do not induce antiviral responses to infection with their endogenous hantavirus, SEOV. This was observed despite high susceptibility to infection and robust viral protein expression, even with low initial MOI. Our investigations further demonstrate that defects in RLR- or type I IFN-dependent signaling in RLMVEC do not explain this immune silence. Finally, we find no evidence of SEOV antagonism of these innate immune signaling pathways. Further work will be necessary to investigate alternative mechanisms of immune evasion such as RNA sequestration, replication efficiency, and metabolic regulation that may play a larger role in the establishment of persistence.

Immune activation, specifically an overactive proinflammatory response, is associated with severe hantavirus disease in humans (69). We have now shown that both HTNV and SEOV drive robust antiviral signaling in human EC, and that this signaling is dependent on the RIG-I-like receptors (**Fig 2** (58)). Results from our RLR^-/-^ HUVEC line indicates that, while HTNV can be detected by both MDA5 and RIG-I, likely in a temporal-specific manner, antiviral responses to SEOV are uniquely RIG-I-dependent. Importantly, induction of ISG expression occurs in HUVEC despite relatively low levels of initial viral infection when compared to Vero E6 cells or RLMVEC (MOI 0.01 vs 0.05 or 0.1, **Fig 1**). This observation suggests that viral PAMPs may be more readily exposed during HUVEC infection, possibly due to poor use of host co-factors required for replication complex formation, mRNA cap-snatching, or complete virion assembly (68). Additionally, this robust RLR-dependent innate immune activation may set the stage for increased infiltration of immune cells, such as proinflammatory neutrophils or monocytes, which have been implicated in severe disease (26, 70).

We were surprised to observe a complete absence of ISG induction in primary RLMVEC following SEOV infection (**Fig 3**). Our experiments were performed with a low starting MOI (0.05) to represent likely physiologic conditions with transmission involving a small number of infectious particles (71). Despite clearly robust SEOV infection on days three and four post-infection, our western blot analysis shows no ISG expression even out to six days post-infection. We would expect that, even if low early levels of infection did not produce a strong IFN signal *in vitro*, by six days post-infection any interferon secretion would drive the amplification of an antiviral response in infected and uninfected cells in culture. Thus, the lack of such a signature is significant. Further, infection with a different strain of SEOV and RLMVEC-passaged SEOV resulted again in an absence of ISG expression. Therefore, we do not believe our observations are a unique artifact of strain history. Infection of rat ECs with HTNV did induce robust ISG expression. Thus, reservoir rat EC mount an antiviral response to non-endemic HTNV infection, but not endemic SEOV. We also interrogated the transcriptional upregulation of *Rig-i, Cxcl10, Ccl2, Mx1*, and *Oas1* by RT-PCR (**Fig 3b, 5a,b**). A scarcity of clean, reliable antibodies for western blotting has limited a deeper investigation for protein expression. However, it is notable that, by assessing both transcription and translation of antiviral genes and ISGs, we have shown that host translational shutoff is not likely to be a mechanism of immune evasion employed by SEOV in its reservoir. The ability of RLMVECs to translate ISGs when infected at high SEOV MOI also supports this conclusion.

Importantly, others have previously investigated antiviral responses by hantavirus reservoir cells to their endemic virus (54, 55, 72). *In vitro* infections of ECs from Norway rats and bank voles have demonstrated that innate immune activation is muted in these cells (54, 55). It, therefore, may be a universal phenotype of hantavirus reservoirs to curb robust innate immune responses to their endemic viruses, representing a potential mechanism of persistence. Dampened immune signaling has been described for several species of bat, reservoirs of highly pathogenic viruses, such as other hemorrhagic fever viruses and rabies virus (7, 9, 73). It is an especially appealing hypothesis to explain the ability of rodents and bats to harbor diverse viral species which otherwise cause immune-mediated pathologies in humans, driven by uncontrolled immune activation. We therefore interrogated the induction of ISGs in response to known agonists of the RLR pathway or type I IFN signaling. Both transfected poly(I:C) and recombinant rat IFNβ drove the transcription of ISGs *Ccl2, Cxcl10*, and *Rig-i* (**Fig 4**). We therefore do not have any indication that the Norwegian rat is an especially immune-privileged host and do not believe this explains the observed lack of ISG expression during SEOV infection.

Given that hantaviruses are thought to have co-evolved along with their rodent reservoir host, we hypothesized that SEOV has evolved mechanisms to directly antagonize antiviral pathways within its natural reservoir to maintain active replication without immune induction (25, 43, 74). We tested whether, in the absence of viral proteins, viral RNAs present during infection are capable of driving innate immune activation. Transfection of total RNA isolated from SEOV-infected Vero E6 cells induced antiviral gene transcription in a dose-dependent manner, suggesting that viral proteins are somehow involved in either antagonizing the RLR pathway or shielding viral RNA from recognition (**Fig 4c**). If SEOV were capable of antagonizing innate immune signaling pathways, we would expect that established SEOV infection would dampen ISG expression driven by type I IFN, poly(I:C), or HTNV. Further, we expected that if SEOV were capable of inhibiting either RIG-I or MDA5 signaling, infection of cells lacking each individual RLR would lead to an inhibition of ISG expression following HTNV infection (**Fig 5**). Importantly, we observed no such inhibition in any of these experiments, providing no evidence to support the hypothesis that SEOV actively antagonizes the RLR or type I IFN signaling cascades in reservoir EC.

A low MOI spreading infection represents a physiologically relevant scenario to study host-pathogen dynamics, with low numbers of infectious particles thought to survive the bottlenecks of host transmission and invasion to establish the initial infection (71, 75). Despite this, it is common for *in vitro* investigations to be performed at very high, non-physiological MOI. Therefore, we investigated whether initiating infection of primary RLMVEC with high SEOV MOI could drive innate immune activation. Indeed, we observed that increasing the initial MOI led to a clear induction of ISGs, with higher MOIs driving earlier gene expression (**Fig 7a**). Notably, even at the highest MOI, ISG expression still does not occur until day 3 post-infection, suggesting that viral propagation and spread is required for immune signaling. This observation is supported by a recent report by the Rockx laboratory, reporting that SEOV infection at an MOI of 1 induced an antiviral transcriptional response in both human and rat ECs (72). Further, we observed that ISG induction in response to high starting MOI was dependent on the RLRs (**Fig 7b**). It is well-established that high MOI infections with negative-sense RNA viruses drives the generation of defective interfering (DI) particles (also referred to as non-standard/defective viral genomes) (65, 67, 68, 76, 77). Recently, a greater appreciation for the role of defective particles and incomplete genomes in triggering innate immune sensors has been highlighted in the literature (78). Thus, the induction of innate immune activation in primary RLMVEC following high MOI infection is possibly due to the accumulation of DI particles. Another possibility is that high MOI infections allow for individual cells to be infected with several viral particles at once, limiting access to critical host factors that facilitate efficient replication and evasion of host PRR, leaving viral PAMPs exposed in the cytosol. Remodeling of intracellular membranes by Tula virus has been proposed as a mechanism of hantavirus sequestration of replication complexes away from surveilling PRR, such as RIG-I and MDA5 (79). SEOV may also be capable of limiting the accumulation of unprocessed RNA or double-stranded RNA intermediates during replication. Trimming of the 5’- triphosphate to a monophosphate on the viral genome is known to be a RIG-I-specific evasion strategy employed by bunyaviruses, including hantaviruses, and may require specific host interactions (80–82). Further research will be necessary to define the mechanism driving immune activation only at high MOI, but our investigations demonstrate the important nuance in interpreting viral data performed only at very high or very low MOI.

We have yet to determine the mechanism(s) by which SEOV limits innate immune activation in low starting MOI conditions, despite strong protein expression and high susceptibility to infection. A deeper understanding of the global SEOV interactome within reservoir and non-reservoir hosts to identify critical protein-protein associations will inform future investigations into these potential host-specific evasion mechanisms. Investigations, like this one, aiming to define the virus-host interactions that allow highly pathogenic zoonotic RNA viruses to be maintained within wild reservoirs are critical to inform public health and wildlife management strategies, as well as provide novel avenues for antiviral therapeutic development.

## Materials and Methods

### Viruses and *in vitro* infections

Seoul virus strain SR11 and Hantaan virus strain 76-118 were propagated on Vero E6 cells (ATCC, CRL-1586) for 12 days, with a maximum of three passages. Infectious virus was isolated by harvesting supernatant and centrifuging at 1000rcf for 10 minutes to remove cellular debris. Rat-passaged SEOV was generated using SR11 stock and propagated over three passages (MOI 0.01) in RLMVECs for ten days each. SEOV Baltimore strain, kindly provided by Dr. Steven Bradfute (UNM), was generated in Vero E6 cells and harvested 12 days post-infection. For virus infections, cells were seeded in cell culture vessels 18-24 hours prior to infection at a target density of 70%. Virus stock was diluted to the target, cell-specific MOI using serum-free Dulbecco’s modified Eagle’s medium (DMEM, VWR 45000-304) supplemented with 1x pen/strep, 1% nonessential amino acids, 2.5% HEPES, and cells were infected for one hour at 37°C. Cells were washed twice with sterile PBS solution (FisherScientific, SH30264FS), and appropriate culture medium was added for the duration of the experiment. For infected cell RNA experiments, Vero E6 cells were infected with SEOV for five days at MOI 0.5 prior to harvest in TRIzol reagent and subsequent RNA extraction. Percent SEOV N positive and infectious unit quantification experiment was performed using SEOV stock virus derived from Vero E6 cells, 1:2 dilution series in serum-free media was performed and cells were infected in a 96-well plate (Agilent, 204624-100) for 1 hour at 37°C followed by addition of methylcellulose (Sigma M0512-500G) overlay with supplemented at 2x concentration DMEM (Gibco, 12100-046) supplemented with 2% FBS, 1x PenStrep, 1% HEPES. UV inactivated virus was treated with UV radiation (FisherBrand UV crosslinker, 13-245-221) until specific absorption of 103.8 mJ/cm^2^.

### Cell Culture

Vero E6 cells (ATCC, CRL-1586) and HEK293T (ATCC, CRL-3216) cells were cultured in Dulbecco’s modified Eagle’s medium (DMEM) supplemented with 10% heat-inactivated FBS, 1% pen/strep, 1% nonessential amino acids, 2.5% HEPES. Primary rat microvascular endothelial cells (RLMVEC, VEC Technologies) were cultured in MCDB-131 base medium (Corning) supplemented with EGMTM-2 Endothelial SingleQuots Bullet Kit (Lonza, CC-4176) and 10% heat-inactivated FBS. Human umbilical endothelial cells (HUVEC-C; ATCC, CRL-1730) were cultured in Lonza EGM-Plus (Lonza, CC-4542) supplemented with bullet kit (Lonza, CC-5036) and 10% heat-inactivated FBS. All endothelial cells were cultured in tissue culture-treated plastics coated with rat tail collagen (VWR, 47747-218). Recombinant rat IFNβ was purchased from R&D Systems (13400–1) and used at 10-150U/milliliter. RLMVEC knockout cells were generated using LentiCrispr V2 plasmid system (83) with gRNAs targeting RIG-I, MDA5, or Cas9 Scramble as control (Table 1). Briefly, HEK293T cells were transfected using ProFection Mammalian Transfection System (Promega, PAE1200) with targeted LentiCrispr V2 plasmid (Addgene Plasmid #98290), pSPAX (Addgene #12260), and p-VSVG (Addgene #138479). One day post-transfection, fresh DMEM was added to HEK293T cells. Virus was harvested two days post-transfection from HEK293T supernatant, filtered through a 0.45μm filter, and added to RLMVECs for 24 hours. Cells were placed under puromycin (VWR, AAJ67236-8EQ) selection at 1μg/milliliter for one week. Knockout efficacy was verified via treatment with rat recombinant IFNβ and immunoblot.

**Table 1.**
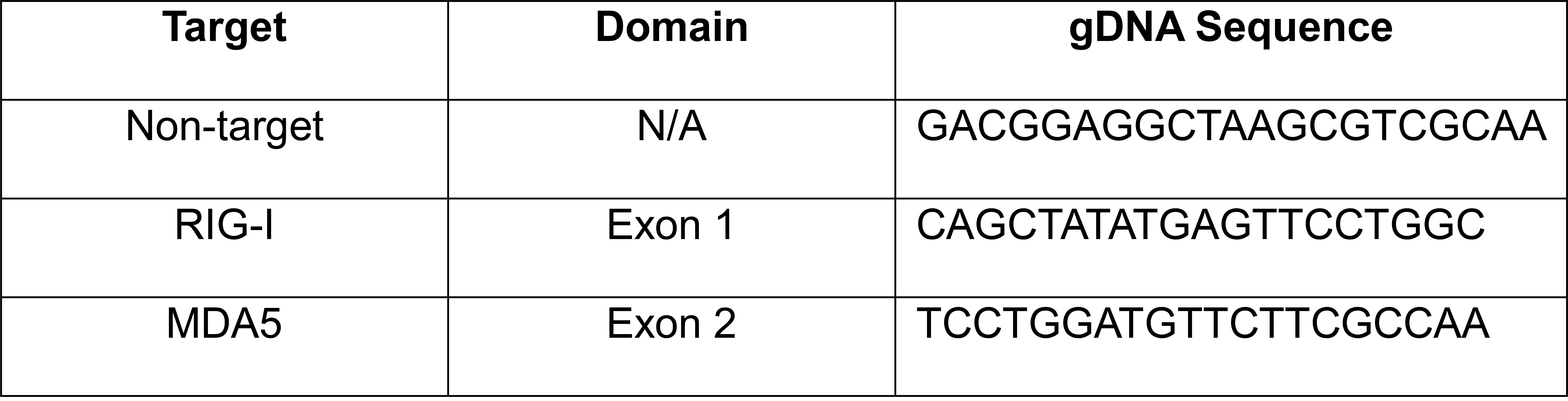
gDNA sequences for RLMVEC CRISPR knockout lines.

### Focus Forming Unit Assay

Infectious virus was quantified using immunostaining as previously described (56). Briefly, Vero E6 cells were infected with 100μL of culture supernatants and incubated for two hours at 37°C. After incubation, a 2% carboxymethylcellulose overlay containing supplemented DMEM (2% pen/strep, 2% nonessential amino acids, 2% HEPES, 4% heat-inactivated FBS). Cells were incubated for seven days at 37°C. After incubation, cells were fixed with 95% EtOH:5% Acetic Acid for 10 minutes at -20°C. Cells were then probed for SEOV N (α SEOV nucleocapsid, custom generated with Genscript) or HTNV N (anti-HTNV nucleocapsid 76–118, BEI resources NR- 12152) at room temperature for 2 hours or 4°C overnight. HRP conjugated secondary antibodies (donkey α rabbit for α HTNV, Jackson Immunoresearch 711-035-152; or goat α mouse for α SEOV, Jackson Immunoresearch 115-035-003) were incubated for 2 hours at room temperature. Foci were then stained using Vector VIP Substrate Kit (Vector Laboratories, SK-4600) and counted under a light microscope to calculate titer.

### Immunofluorescence and Microscopy Analysis

To calculate SEOV N positivity and cell-specific titers, Vero E6, HUVEC, and RLMVEC were cultured in 96-well plates and infected with serial dilutions of SEOV as described in infection methods above. 24 hours post-infection, cells were fixed with 95% EtOH:5% Acetic Acid solution for ten minutes at -20°C then blocked for one hour in 3% FCS in PBS. Cells were probed overnight at 4°C for SEOV N protein (custom, Genscript) at dilution 1:400 in 1xPBS and with secondary antibody AlexaFluor 555 goat α mouse (Thermo Fisher Scientific, A-31570) at dilution 1:400 for two hours at room temperature. Nuclear stain DAPI was used at 1:1000 for ten minutes at room temperature (SeraCare, 5930-0006). High-content imaging was acquired on CellInsight CX7 High-content Analysis platform (Thermo Fisher Scienitific, CX7A1110) at 10x objective magnification using iDev software. SEOV N positive cells were quantified using the Spot Detector Applet in the iDev software. Regions of interest (ROI) were defined based on DAPI nuclear stain, and SEOV N positive cells were defined as having at least one spot within the ROI as thresholded relative to each cell type. Data were collected on a total of 25 fields per well for all cell types. Representative images were acquired on the EVOS FL Auto imaging system (Thermo Fisher Scientific, AMF7000). RLMVECs assayed for Mx1/2/3 expression during SEOV infection were fixed using 4% paraformaldehyde (VWR, 97064-606) solution in sterile PBS for 30 minutes at room temperature, then permeabilized with 0.01% Triton X-100 (VWR, 97062-208) and blocked for one hour in 3% FCS in PBS. Cells were treated with DAPI nuclear stain 1:1000, SEOV N nucleocapsid 1:400, or Mx 1/2/3 1:400 in PBS (Santa Cruz Biotechnology, sc-166412 AF488). Representative images were acquired via the EVOS FL Auto imaging system (Thermo Fisher Scientific).

### Cell-specific Titer Calculations

Cell-specific titers were calculated by determining the number of SEOV N-expressing cells for each cell type using a serial dilution of SEOV stock. This method is adapted from a recent publication by Menke et al. (57). Briefly, total number of N expressing cells was quantified as described above. For each dilution:

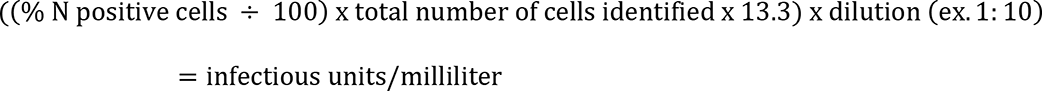

Titer was calculated for all replicate wells in each dilution for which the number of N-positive cells decreased two-fold, to match the inoculum dilutions. Replicates were averaged and then the calculated titers for each dilution condition were averaged to determine the titer on each cell type. Four replicates per dilution and >3 independent experiments were performed for each cell type.

### RNA Methods

Cells were lysed for RNA analysis using TRIzol Reagent (ThermoFisher Scientific, 15596026). RNA used for infected cell RNA transfection was isolated using phenol:chloroform extraction following manufacturer protocol. RNA used for gene expression analysis was purified using Zymo Research Direct-zol RNA Miniprep Plus kit (VWR 76211-340) according to manufacturer’s instructions. RNA concentrations were quantified using a NanoDrop UV-Vis Spectrophotometer (ND-1000). cDNA was synthesized using Applied Biosystems High-Capacity cDNA Reverse Transcription Kit (FisherScientific, 43-688-14) by adding 400ng total RNA to the reaction mixture containing random primers following manufacturer guidelines. Real-time PCR was performed using the SYBR Green PCR Master Mix (ThermoFisher Scientific, 4364346) and the primers described in Table 1. In-house designed primers were purchased from Integrated DNA Technologies and TaqMan primer assays were purchased from ThermoFisher. Host antiviral ISG expression was normalized to rat Chmp2a mRNA, and fold change calculated as ddCt over mock infection. Results were visualized and statistically analyzed using GraphPad Prism software (Prism 10). Synthetic polyinosinic:polycytidylic acid (poly(I:C)) was purchased from Sigma-Aldrich (MilliporeSigma, P0913-50MG). The hepatitis C virus (HCV) xRNA was *in vitro* transcribed from synthetic DNA oligonucleotide templates (Integrated DNA Technologies) using the T7 MEGAshortscript kit (Ambion) as previously described (84–86). Following manufacturer protocol, transfections were completed using JetPRIME transfection reagent (VWR, 89129-924).

### Protein Methods

Cell lysates to be analyzed for protein expression by immunoblot analysis were harvested in protein lysis buffer and prepared as previously described (87). Briefly, protein was harvested in RIPA buffer and clarified through 25,000rcf centrifugation for 15 minutes at 4°C. Protein was quantified via Pierce BCA Protein Assay Kit (ThermoFisher Scientific, 23225). 15μg-30ug of protein was loaded per sample into a 10% polyacrylamide gel and, after denaturing electrophoresis, transferred to a 0.45µm nitrocellulose membrane (VWR, 10120-006). Membranes were blocked at room temperature in 10% FCS in PBS-T. Primary antibodies (table 2) were incubated at 4° overnight. HRP-conjugated secondary antibodies against primary antibody species were incubated for 1 hour at room temperature. Blots were imaged on BioRad Chemidoc MP Imaging System using chemiluminescence Pierce Substrate for Western Blotting (VWR, PI80196).

**Table 2.**
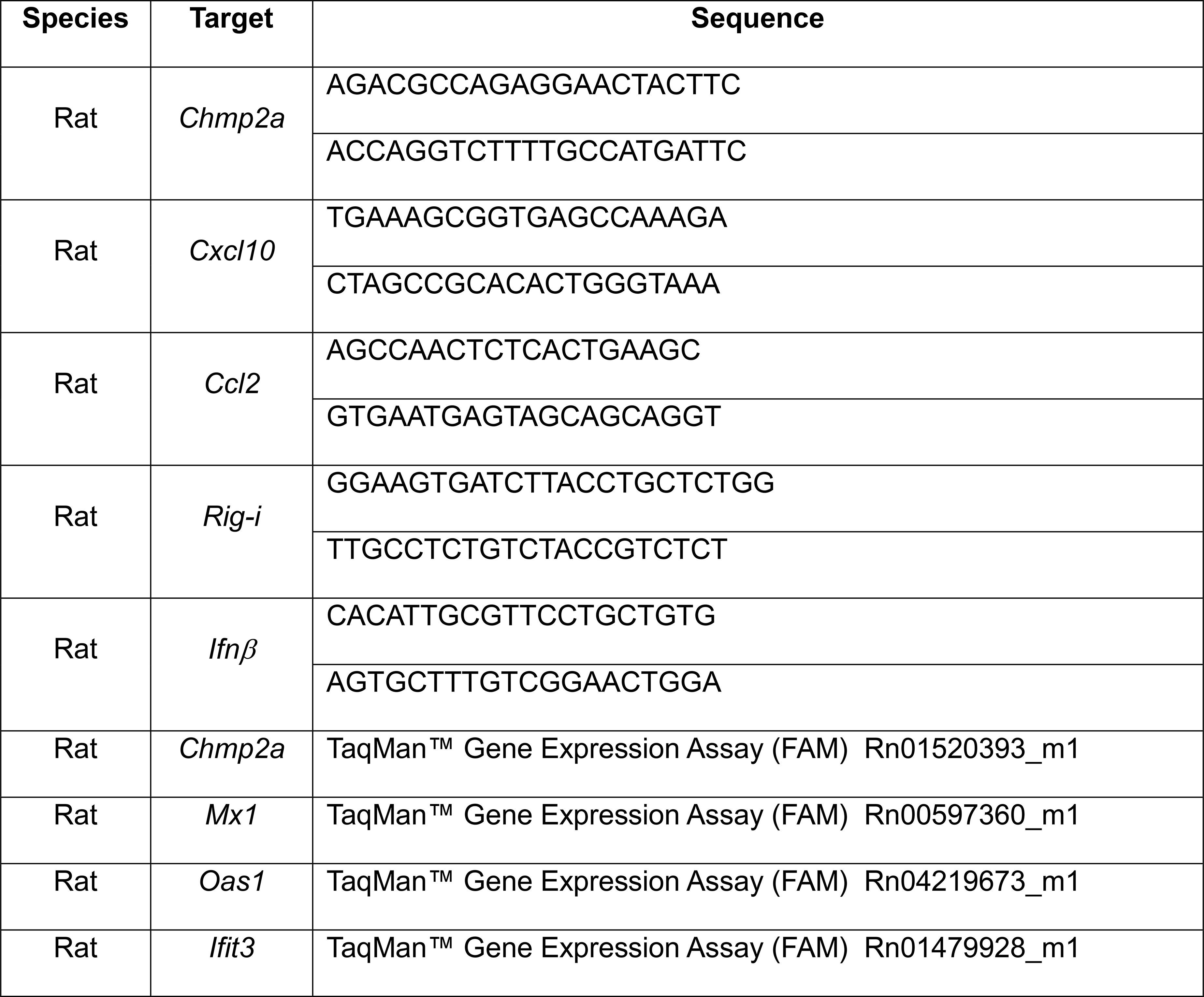
Primer sequences or catalog numbers for real-time PCR analyses.

**Table 3.** Primary antibodies used for protein detection in immunoblot and immunofluorescence assays.

| Target Species | Protein | Manufacturer |
| --- | --- | --- |
| Rat | Mx 1/2/3 | SantaCruz Biotech (sc-166412) |
| Rat | Mx 1/2/3 AF488 | SantaCruz Biotech (sc-166412 AF488) |
| Rat | MDA5 | GeneTex (GTX103138) |
| Rat | RIG-I | Cell Signaling Technologies (3743T) |
| Human, Rat | βActin | SantaCruz Biotech (sc-47778) |
| Human | Mx | Cell Signaling (37849S) |
| Human | IFIT1 | Abcam (ab236256) |
| SEOV | SEOV Nucleocapsid | Custom, Genscript |

### Secreted cytokine analysis

Tissue culture supernatant from HUVEC (Cas9 and RLR^-/-^) was collected four days post SEOV infection (MOI 0.01) and clarified for cellular debris by centrifugation (1000 rcf for 10 minutes). Clarified supernatants were UV-inactivated (specific absorption of 103.8 mJ/cm^2^) and shipped to Eve Technologies for interrogation on the Human Cytokine/Chemokine Panel A 48-Plex Discovery Assay® Array (HD48A). One experiment was performed with three biological replicates and two technical assay replicates.

### Statistical analyses

Statistical significance was assess using GraphPad Prism software version 10. Where noted in figure legends, ANOVA and student’s T-test analyses were applied, with * representing adjusted p<0.05. All comparisons reaching statistical significance are displayed on graphs.

## Acknowledgements

We would like to acknowledge Dr. Sharina Desai and the Autophagy, Inflammation, and Metabolism core for resource support and technical expertise. We also acknowledge Dr. Steven Bradfute and Dr. Curt Hines for resource sharing and constructive feedback on this project. We thank Dr. Adriana Forero and Dr. Emily Hemann for helpful critiques on this manuscript.

## References

1. Allen LJ, McCormack RK, Jonsson CB. Mathematical models for hantavirus infection in rodents. Bulletin of mathematical biology. 2006;68(3):511–24.

2. Allen LJ, Wesley CL, Owen RD, Goodin DG, Koch D, Jonsson CB, et al. A habitat-based model for the spread of hantavirus between reservoir and spillover species. Journal of theoretical biology. 2009;260(4):510–22.

3. Jonsson CB, Figueiredo LT, Vapalahti O. A global perspective on hantavirus ecology, epidemiology, and disease. Clin Microbiol Rev. 2010;23(2):412–41.

4. Han BA, Schmidt JP, Bowden SE, Drake JM. Rodent reservoirs of future zoonotic diseases. Proc Natl Acad Sci U S A. 2015;112(22):7039–44.

5. Barbour AG. Infection resistance and tolerance in Peromyscus spp., natural reservoirs of microbes that are virulent for humans. Semin Cell Dev Biol. 2017;61:115–22.

6. Balderrama-Gutierrez G, Milovic A, Cook VJ, Islam MN, Zhang Y, Kiaris H, et al. An Infection-Tolerant Mammalian Reservoir for Several Zoonotic Agents Broadly Counters the Inflammatory Effects of Endotoxin. mBio. 2021;12(2).

7. Cockrell C, An G. Comparative Computational Modeling of the Bat and Human Immune Response to Viral Infection with the Comparative Biology Immune Agent Based Model. Viruses. 2021;13(8).

8. Silvestri G. Naturally SIV-infected sooty mangabeys: are we closer to understanding why they do not develop AIDS? J Med Primatol. 2005;34(5-6):243–52.

9. Skirmuntt EC, Escalera-Zamudio M, Teeling EC, Smith A, Katzourakis A. The Potential Role of Endogenous Viral Elements in the Evolution of Bats as Reservoirs for Zoonotic Viruses. Annu Rev Virol. 2020;7(1):103–19.

10. Eby P, Peel AJ, Hoegh A, Madden W, Giles JR, Hudson PJ, et al. Pathogen spillover driven by rapid changes in bat ecology. Nature. 2023;613(7943):340-4.

11. Mandl JN, Ahmed R, Barreiro LB, Daszak P, Epstein JH, Virgin HW, et al. Reservoir host immune responses to emerging zoonotic viruses. Cell. 2015;160(1-2):20–35.

12. Brook CE, Rozins C, Guth S, Boots M. Reservoir host immunology and life history shape virulence evolution in zoonotic viruses. PLoS Biol. 2023;21(9):e3002268.

13. Seal S, Dharmarajan G, Khan I. Evolution of pathogen tolerance and emerging infections: A missing experimental paradigm. Elife. 2021;10.

14. Dowling JW, Forero A. Beyond Good and Evil: Molecular Mechanisms of Type I and III IFN Functions. J Immunol. 2022;208(2):247–56.

15. Brisse M, Ly H. Comparative Structure and Function Analysis of the RIG-I-Like Receptors: RIG-I and MDA5. Front Immunol. 2019;10:1586.

16. Chan YK, Gack MU. RIG-I-like receptor regulation in virus infection and immunity. Curr Opin Virol. 2015;12:7–14.

17. Kell AM, Gale M, Jr. RIG-I in RNA virus recognition. Virology. 2015;479–480:110-21.

18. Capone F, Guerriero E, Colonna G, Maio P, Mangia A, Castello G, et al. Cytokinome profile evaluation in patients with hepatitis C virus infection. World J Gastroenterol. 2014;20(28):9261–9.

19. Gu Y, Hsu AC, Pang Z, Pan H, Zuo X, Wang G, et al. Role of the Innate Cytokine Storm Induced by the Influenza A Virus. Viral Immunol. 2019;32(6):244–51.

20. Creisher PS, Klein SL. Pathogenesis of viral infections during pregnancy. Clin Microbiol Rev. 2024:e0007323.

21. Batiha GE, Al-Kuraishy HM, Al-Gareeb AI, Welson NN. Pathophysiology of Post-COVID syndromes: a new perspective. Virol J. 2022;19(1):158.

22. DePaula-Silva AB. The Contribution of Microglia and Brain-Infiltrating Macrophages to the Pathogenesis of Neuroinflammatory and Neurodegenerative Diseases during TMEV Infection of the Central Nervous System. Viruses. 2024;16(1).

23. Botten J, Mirowsky K, Kusewitt D, Ye C, Gottlieb K, Prescott J, et al. Persistent Sin Nombre virus infection in the deer mouse (Peromyscus maniculatus) model: sites of replication and strand-specific expression. J Virol. 2003;77(2):1540–50.

24. Chu YK, Owen RD, Gonzalez LM, Jonsson CB. The complex ecology of hantavirus in Paraguay. The American journal of tropical medicine and hygiene. 2003;69(3):263–8.

25. Guo WP, Lin XD, Wang W, Tian JH, Cong ML, Zhang HL, et al. Phylogeny and origins of hantaviruses harbored by bats, insectivores, and rodents. PLoS pathogens. 2013;9(2):e1003159.

26. Strandin T, Mäkelä S, Mustonen J, Vaheri A. Neutrophil Activation in Acute Hemorrhagic Fever With Renal Syndrome Is Mediated by Hantavirus-Infected Microvascular Endothelial Cells. Front Immunol. 2018;9:2098.

27. Angulo J, Martinez-Valdebenito C, Marco C, Galeno H, Villagra E, Vera L, et al. Serum levels of interleukin-6 are linked to the severity of the disease caused by Andes Virus. PLoS neglected tropical diseases. 2017;11(7):e0005757.

28. Bondu V, Schrader R, Gawinowicz MA, McGuire P, Lawrence DA, Hjelle B, et al. Elevated cytokines, thrombin and PAI-1 in severe HCPS patients due to Sin Nombre virus. Viruses. 2015;7(2):559–89.

29. Koskela S, Mäkelä S, Strandin T, Vaheri A, Outinen T, Joutsi-Korhonen L, et al. Coagulopathy in Acute Puumala Hantavirus Infection. Viruses. 2021;13(8).

30. Solà-Riera C, Gupta S, Maleki KT, González-Rodriguez P, Saidi D, Zimmer CL, et al. Hantavirus Inhibits TRAIL-Mediated Killing of Infected Cells by Downregulating Death Receptor 5. Cell Rep. 2019;28(8):2124–39.e6.

31. Solà-Riera C, Gupta S, Ljunggren HG, Klingström J. Orthohantaviruses belonging to three phylogroups all inhibit apoptosis in infected target cells. Sci Rep. 2019;9(1):834.

32. Solà-Riera C, García M, Ljunggren HG, Klingström J. Hantavirus inhibits apoptosis by preventing mitochondrial membrane potential loss through up-regulation of the pro-survival factor BCL-2. PLoS Pathog. 2020;16(2):e1008297.

33. Vaheri A, Strandin T, Hepojoki J, Sironen T, Henttonen H, Makela S, et al. Uncovering the mysteries of hantavirus infections. Nature reviews Microbiology. 2013;11(8):539–50.

34. Kell AM, Hemann EA, Turnbull JB, Gale M. RIG-I-like receptor activation drives type I IFN and antiviral signaling to limit Hantaan orthohantavirus replication. PLoS Pathog. 2020;16(4):e1008483.

35. Martínez VP, Di Paola N, Alonso DO, Pérez-Sautu U, Bellomo CM, Iglesias AA, et al. “Super-Spreaders” and Person-to-Person Transmission of Andes Virus in Argentina. N Engl J Med. 2020;383(23):2230–41.

36. Peters CJ, Khan AS. Hantavirus pulmonary syndrome: the new American hemorrhagic fever. Clin Infect Dis. 2002;34(9):1224–31.

37. Guo J, Guo X, Wang Y, Tian F, Luo W, Zou Y. Cytokine response to Hantaan virus infection in patients with hemorrhagic fever with renal syndrome. Journal of medical virology. 2017;89(7):1139–45.

38. Rasmuson J, Pourazar J, Mohamed N, Lejon K, Evander M, Blomberg A, et al. Cytotoxic immune responses in the lungs correlate to disease severity in patients with hantavirus infection. European journal of clinical microbiology & infectious diseases : official publication of the European Society of Clinical Microbiology. 2016;35(4):713–21.

39. Botten J, Mirowsky K, Kusewitt D, Bharadwaj M, Yee J, Ricci R, et al. Experimental infection model for Sin Nombre hantavirus in the deer mouse (Peromyscus maniculatus). Proc Natl Acad Sci U S A. 2000;97(19):10578–83.

40. Campbell CL, Torres-Perez F, Acuna-Retamar M, Schountz T. Transcriptome markers of viral persistence in naturally-infected andes virus (bunyaviridae) seropositive long-tailed pygmy rice rats. PloS one. 2015;10(4):e0122935.

41. Easterbrook JD, Zink MC, Klein SL. Regulatory T cells enhance persistence of the zoonotic pathogen Seoul virus in its reservoir host. Proceedings of the National Academy of Sciences of the United States of America. 2007;104(39):15502–7.

42. Easterbrook JD, Klein SL. Immunological mechanisms mediating hantavirus persistence in rodent reservoirs. PLoS pathogens. 2008;4(11):e1000172.

43. Holmes EC, Zhang YZ. The evolution and emergence of hantaviruses. Current opinion in virology. 2015;10:27–33.

44. Compton SR, Jacoby RO, Paturzo FX, Smith AL. Persistent Seoul virus infection in Lewis rats. Arch Virol. 2004;149(7):1325–39.

45. Easterbrook JD, Shields T, Klein SL, Glass GE. Norway rat population in Baltimore, Maryland, 2004. Vector borne and zoonotic diseases. 2005;5(3):296-9.

46. Griffiths J, Yeo HL, Yap G, Mailepessov D, Johansson P, Low HT, et al. Survey of rodent- borne pathogens in Singapore reveals the circulation of Leptospira spp., Seoul hantavirus, and Rickettsia typhi. Sci Rep. 2022;12(1):2692.

47. He W, Fu J, Wen Y, Cheng M, Mo Y, Chen Q. Detection and Genetic Characterization of Seoul Virus in Liver Tissue Samples From. Front Vet Sci. 2021;8:748232.

48. Kerins JL, Koske SE, Kazmierczak J, Austin C, Gowdy K, Dibernardo A. Outbreak of Seoul Virus Among Rats and Rat Owners - United States and Canada, 2017. MMWR Morbidity and mortality weekly report. 2018;67(4):131-4.

49. Knust B, Brown S, de St Maurice A, Whitmer S, Koske SE, Ervin E, et al. Seoul Virus Infection and Spread in United States Home-Based Ratteries: Rat and Human Testing Results From a Multistate Outbreak Investigation. J Infect Dis. 2020;222(8):1311–9.

50. Green W, Feddersen R, Yousef O, Behr M, Smith K, Nestler J, et al. Tissue distribution of hantavirus antigen in naturally infected humans and deer mice. J Infect Dis. 1998;177(6):1696–700.

51. Lehmer EM, Jones JD, Bego MG, Varner JM, Jeor SS, Clay CA, et al. Long-term patterns of immune investment by wild deer mice infected with Sin Nombre virus. Physiol Biochem Zool. 2010;83(5):847–57.

52. Schountz T, Quackenbush S, Rovnak J, Haddock E, Black WCt, Feldmann H, et al. Differential lymphocyte and antibody responses in deer mice infected with Sin Nombre hantavirus or Andes hantavirus. Journal of virology. 2014;88(15):8319–31.

53. Spengler JR, Haddock E, Gardner D, Hjelle B, Feldmann H, Prescott J. Experimental Andes virus infection in deer mice: characteristics of infection and clearance in a heterologous rodent host. PLoS One. 2013;8(1):e55310.

54. Li W, Klein SL. Seoul virus-infected rat lung endothelial cells and alveolar macrophages differ in their ability to support virus replication and induce regulatory T cell phenotypes. Journal of virology. 2012;86(21):11845–55.

55. Strandin T, Smura T, Ahola P, Aaltonen K, Sironen T, Hepojoki J, et al. Orthohantavirus Isolated in Reservoir Host Cells Displays Minimal Genetic Changes and Retains Wild-Type Infection Properties. Viruses. 2020;12(4).

56. Kraus AA, Priemer C, Heider H, Kruger DH, Ulrich R. Inactivation of Hantaan virus- containing samples for subsequent investigations outside biosafety level 3 facilities. Intervirology. 2005;48(4):255–61.

57. Menke L, Sieben C. An Improved Workflow for the Quantification of Orthohantavirus Infection Using Automated Imaging and Flow Cytometry. Viruses. 2024;16(2).

58. Kell AM, Hemann EA, Turnbull JB, Gale M, Jr. RIG-I-like receptor activation drives type I IFN and antiviral signaling to limit Hantaan orthohantavirus replication. PLoS Pathog. 2020;16(4):e1008483.

59. Xiao SY, Leduc JW, Chu YK, Schmaljohn CS. Phylogenetic analyses of virus isolates in the genus Hantavirus, family Bunyaviridae. Virology. 1994;198(1):205–17.

60. Prescott J, Feldmann H, Safronetz D. Amending Koch’s postulates for viral disease: When “growth in pure culture” leads to a loss of virulence. Antiviral Res. 2017;137:1–5.

61. Lundkvist A, Cheng Y, Sjolander KB, Niklasson B, Vaheri A, Plyusnin A. Cell culture adaptation of Puumala hantavirus changes the infectivity for its natural reservoir, Clethrionomys glareolus, and leads to accumulation of mutants with altered genomic RNA S segment. Journal of virology. 1997;71(12):9515–23.

62. You S, Rice CM. 3’ RNA elements in hepatitis C virus replication: kissing partners and long poly(U). Journal of virology. 2008;82(1):184–95.

63. Saito T, Owen DM, Jiang F, Marcotrigiano J, Gale M, Jr. Innate immunity induced by composition-dependent RIG-I recognition of hepatitis C virus RNA. Nature. 2008;454(7203):523- 7.

64. Marshall-Clarke S, Downes JE, Haga IR, Bowie AG, Borrow P, Pennock JL, et al. Polyinosinic acid is a ligand for toll-like receptor 3. J Biol Chem. 2007;282(34):24759–66.

65. Mercado-Lopez X, Cotter CR, Kim WK, Sun Y, Munoz L, Tapia K, et al. Highly immunostimulatory RNA derived from a Sendai virus defective viral genome. Vaccine. 2013;31(48):5713–21.

66. Xu J, Mercado-Lopez X, Grier JT, Kim WK, Chun LF, Irvine EB, et al. Identification of a Natural Viral RNA Motif That Optimizes Sensing of Viral RNA by RIG-I. mBio. 2015;6(5):e01265–15.

67. Ziegler CM, Botten JW. Defective Interfering Particles of Negative-Strand RNA Viruses. Trends Microbiol. 2020;28(7):554–65.

68. Vignuzzi M, López CB. Defective viral genomes are key drivers of the virus-host interaction. Nat Microbiol. 2019;4(7):1075–87.

69. Wang PZ, Li ZD, Yu HT, Zhang Y, Wang W, Jiang W, et al. Elevated serum concentrations of inflammatory cytokines and chemokines in patients with haemorrhagic fever with renal syndrome. The Journal of international medical research. 2012;40(2):648–56.

70. Koma T, Yoshimatsu K, Nagata N, Sato Y, Shimizu K, Yasuda SP, et al. Neutrophil depletion suppresses pulmonary vascular hyperpermeability and occurrence of pulmonary edema caused by hantavirus infection in C.B-17 SCID mice. Journal of virology. 2014;88(13):7178-88.

71. Gutiérrez S, Michalakis Y, Blanc S. Virus population bottlenecks during within-host progression and host-to-host transmission. Curr Opin Virol. 2012;2(5):546–55.

72. Noack D, van den Hout MCGN, Embregts CWE, van IJcken WFJ, Koopmans MPG, Rockx B. Species-specific responses during Seoul orthohantavirus infection in human and rat lung microvascular endothelial cells. PLoS Negl Trop Dis. 2024;18(3):e0012074.

73. Ahn M, Chen VC, Rozario P, Ng WL, Kong PS, Sia WR, et al. Bat ASC2 suppresses inflammasomes and ameliorates inflammatory diseases. Cell. 2023;186(10):2144–59.e22.

74. Plyusnin A, Sironen T. Evolution of hantaviruses: co-speciation with reservoir hosts for more than 100 MYR. Virus Res. 2014;187:22–6.

75. Botten J, Mirowsky K, Ye C, Gottlieb K, Saavedra M, Ponce L, et al. Shedding and intracage transmission of Sin Nombre hantavirus in the deer mouse (Peromyscus maniculatus) model. J Virol. 2002;76(15):7587–94.

76. Ziegler CM, Eisenhauer P, Bruce EA, Weir ME, King BR, Klaus JP, et al. The Lymphocytic Choriomeningitis Virus Matrix Protein PPXY Late Domain Drives the Production of Defective Interfering Particles. PLoS Pathog. 2016;12(3):e1005501.

77. Manzoni TB, López CB. Defective (interfering) viral genomes re-explored: impact on antiviral immunity and virus persistence. Future Virol. 2018;13(7):493–503.

78. Kagan JC. Infection infidelities drive innate immunity. Science. 2023;379(6630):333-5.

79. Davies KA, Chadwick B, Hewson R, Fontana J, Mankouri J, Barr JN. The RNA Replication Site of Tula Orthohantavirus Resides within a Remodelled Golgi Network. Cells. 2020;9(7).

80. Garcin D, Lezzi M, Dobbs M, Elliott RM, Schmaljohn C, Kang CY, et al. The 5’ ends of Hantaan virus (Bunyaviridae) RNAs suggest a prime-and-realign mechanism for the initiation of RNA synthesis. J Virol. 1995;69(9):5754–62.

81. Habjan M, Andersson I, Klingstrom J, Schumann M, Martin A, Zimmermann P, et al. Processing of genome 5’ termini as a strategy of negative-strand RNA viruses to avoid RIG-I- dependent interferon induction. PLoS One. 2008;3(4):e2032.

82. Hornung V, Ellegast J, Kim S, Brzozka K, Jung A, Kato H, et al. 5’-Triphosphate RNA is the ligand for RIG-I. Science. 2006;314(5801):994-7.

83. Sanjana NE, Shalem O, Zhang F. Improved vectors and genome-wide libraries for CRISPR screening. Nat Methods. 2014;11(8):783–4.

84. Saito T, Gale M, Jr. Differential recognition of double-stranded RNA by RIG-I-like receptors in antiviral immunity. The Journal of experimental medicine. 2008;205(7):1523–7.

85. Schnell G, Loo YM, Marcotrigiano J, Gale M, Jr. Uridine composition of the poly-U/UC tract of HCV RNA defines non-self recognition by RIG-I. PLoS pathogens. 2012;8(8):e1002839.

86. Kell A, Stoddard M, Li H, Marcotrigiano J, Shaw GM, Gale M, Jr. Pathogen-Associated Molecular Pattern Recognition of Hepatitis C Virus Transmitted/Founder Variants by RIG-I Is Dependent on U-Core Length. J Virol. 2015;89(21):11056–68.

87. Foy E, Li K, Sumpter R, Jr., Loo YM, Johnson CL, Wang C, et al. Control of antiviral defenses through hepatitis C virus disruption of retinoic acid-inducible gene-I signaling. Proceedings of the National Academy of Sciences of the United States of America. 2005;102(8):2986–91.

